# Genetic architecture drives seasonal onset of hibernation in the 13-lined ground squirrel

**DOI:** 10.1101/222307

**Authors:** Katharine R. Grabek, Thomas F. Cooke, L. Elaine Epperson, Kaitlyn K. Spees, Gleyce F. Cabral, Shirley C. Sutton, Dana K. Merriman, Sandy L. Martin, Carlos D. Bustamante

## Abstract

Hibernation is a highly dynamic phenotype whose timing, for many mammals, is controlled by a circannual clock and accompanied by rhythms in body mass and food intake. When housed in an animal facility, 13-lined ground squirrels exhibit individual variation in the seasonal onset of hibernation, which is not explained by environmental or biological factors, such as body mass and sex. We hypothesized that underlying genetic architecture instead drives variation in this timing. After first increasing the contiguity of the genome assembly, we therefore employed a genotype-by-sequencing approach to characterize genetic variation in 153 13-lined ground squirrels. Combining this with datalogger records, we estimated high heritability (61-100%) for the seasonal onset of hibernation. After applying a genome-wide scan with 46,996 variants, we also identified 21 loci significantly associated with hibernation immergence, which alone accounted for 54% of the variance in the phenotype. The most significant marker (SNP 15, p=3.81×10−6) was located near *prolactin*-*releasing hormone receptor* (*PRLHR*), a gene that regulates food intake and energy homeostasis. Other significant loci were located near genes functionally related to hibernation physiology, including *muscarinic acetylcholine receptor M2* (*CHRM2*), involved in the control of heart rate, *exocyst complex component 4* (*EXOC4*) *and prohormone convertase 2* (*PCSK2*), both of which are involved in insulin signaling and processing. Finally, we applied an expression quantitative loci (eQTL) analysis using existing transcriptome datasets, and we identified significant (q<0.1) associations for 9/21 variants. Our results highlight the power of applying a genetic mapping strategy to hibernation and present new insight into the genetics driving its seasonal onset.

## Introduction

Hibernation is a highly dynamic phenotype that maximizes energy savings during periods of low resource availability. For a number of mammals, such as the 13-lined ground squirrel, *Ictidomys tridecemlineatus,* an endogenous circannual clock controls the timing of winter hibernation, along with rhythms in reproductive behavior, body mass, and food intake [1-3]. These hibernators partition their year between two distinct states, homeothermy and heterothermy (a.k.a hibernation, Fig 1A) that are distinguished by dramatic differences in behavior and physiology. While physiology during homeothermy resembles that of a nonhibernating mammal, squirrels spend most of their hibernation time in an energy-conserving state called torpor (Fig 1B, top right). Here, metabolic, respiratory and heart rates are dramatically reduced to 1-9% of homeothermic baselines, while body temperature is lowered to near freezing [4]. However, torpor is not continuous, but instead punctuated by brief, metabolically intense, arousals that largely restore baseline physiology, including near-homeothermic body temperature [5,6]. Thus, hibernation is a period of heterothermy composed of cycles between torpor and arousal.

**Fig 1.**
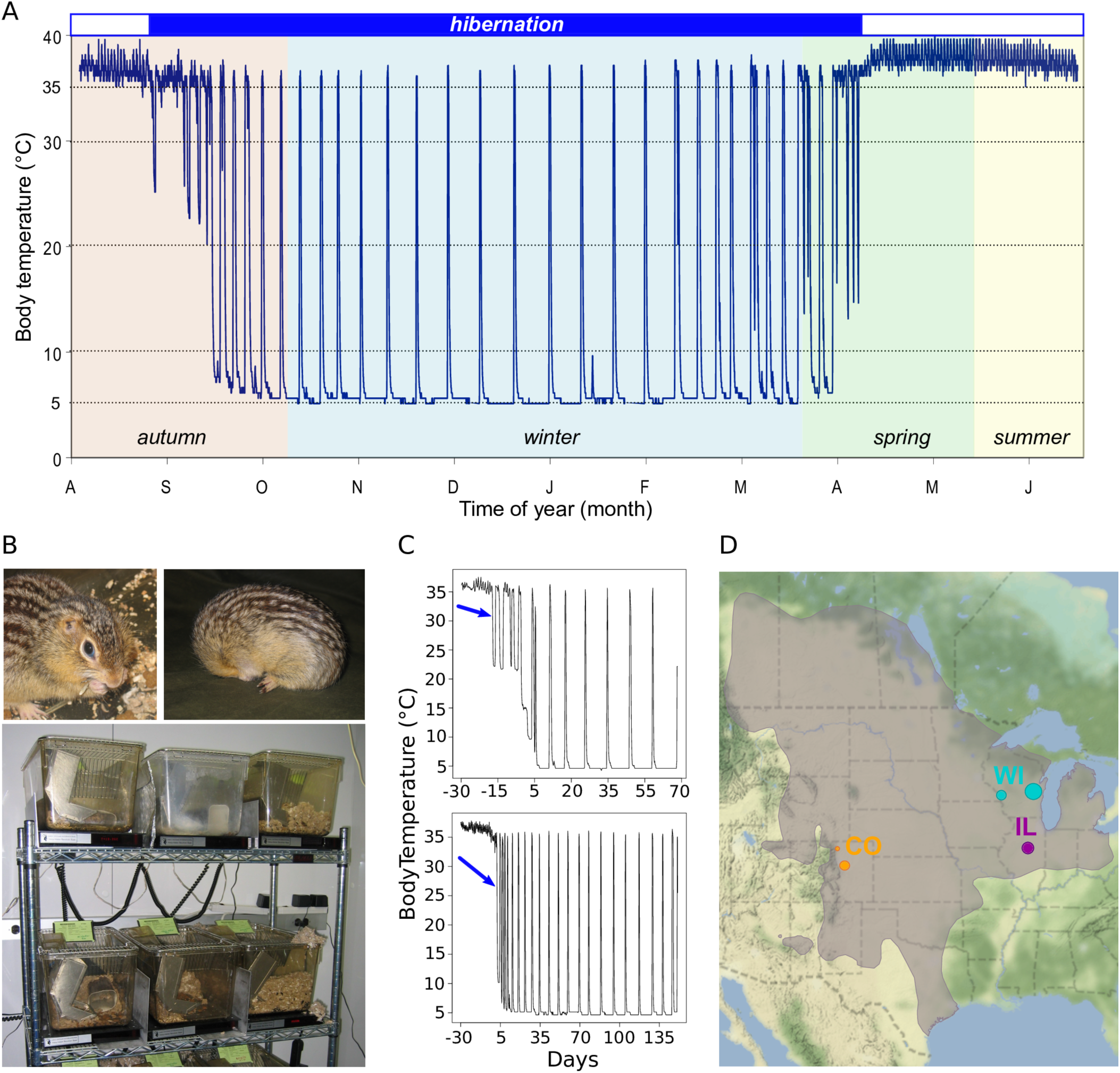
The 13-lined ground squirrel as a model for studying the genetics of hibernation. (A) Body temperature trace showing a 13-lined ground squirrel’s year. Hibernating portion is demarcated by blue shaded box above. (B) A non-hibernating (top left) and hibernating (top right) 13-lined ground squirrel, individually housed in standard lab rodent cages in an animal facility (bottom). (C) Representative plots of body temperature telemetry analyses. Arrows point to the first day of torpor, the phenotype measured in this study. Days are ± from hibernaculum placement. (D) Approximate locales of squirrels genotyped in this study. Shaded area indicates the 13-lined ground squirrel’s geographic range [102].

The seasonal transition from homeothermy to heterothermy occurs during the autumn of each year. Successful hibernation requires preparation, most notably the storage of large amounts of energy in the form of fat, because this species fasts throughout the heterothermic period. While post-reproduction homeothermy is marked by increased food intake, as the onset of heterothermy approaches, the squirrel’s metabolic rate slows, peak body mass is achieved, and food intake ceases [7]. At the cellular level, glucose-based metabolism is switched to one that is primarily lipid-based, and lipogenesis is swapped for lipolysis [8]. While peak plasma insulin concentration occurs during this period, paradoxically, animals also become transiently insulin-resistant [9]. At the mRNA and protein levels, sweeping changes in expression are observed [10-14]. However, the genetic factors driving this transition remain largely unknown.

The commencement of torpor, defined by a criterion drop in body temperature, is one readily quantifiable outcome of the transition that marks the start of seasonal heterothermy. When housed under standard laboratory conditions in an animal facility (Fig 1B, bottom), 13-lined ground squirrels exhibit individual variation in the timing of their first bout of torpor. This variation is neither accounted for by environmental signals, such as food withdrawal, shortened photoperiod, or falling ambient temperature, nor by biological factors, such as age, body mass, and sex. All of these variables have little to modest influence on timing [15]. Rather, consistent with being controlled by an endogenous circannual clock, we hypothesized that observed variation in the onset of torpor is due to underlying genetic variation between individuals. If so, applying a genome-wide scan could potentially identify genetic components driving the start of seasonal heterothermy.

Therefore, in this study, we first increased the contiguity of the 13-lined ground squirrel draft genome assembly. We next employed a genotype-by-sequencing strategy to characterize genetic variation in 153 13-lined ground squirrels whose tissues were previously collected for use in transcriptomic, proteomic and biochemical studies [16-22]. Many of these squirrels were surgically implanted with body temperature dataloggers, and from their records, we recorded the first day that torpor occurred in each individual (Fig 1C). We next estimated the heritability of, and identified genetic variants associated with, the onset of autumn torpor in this species. Finally, we integrated data from prior transcriptomic studies to identify transcripts whose expression levels were significantly associated with these variants. Our results present new insight into the genetics driving the transition from homeothermy to heterothermy and illustrate the power of genetic analysis to attack questions of exceptional biological significance in a non-classical genetic model organism.

## Results

### Long-range scaffolding of the draft genome assembly

At the time this study began, the existing 13-lined ground squirrel genome assembly (like that of many non-model organisms) contained thousands of unordered scaffolds, which could lead to difficulties in identifying causative variants, as peaks in linkage disequilibrium (LD) could be spread across multiple scaffolds. We therefore first sought to increase the genome’s contiguity using a long-range scaffolding technique [23].

A single library was constructed using proximity ligation of in vitro reconstituted chromatin. After sequencing, which provided 52.6x physical coverage of the genome (Table 1) and scaffolding, the contiguity of the final HiRise assembly was increased approximately three-fold as compared to the existing draft assembly (N50 of 22.6Mb vs 8.19Mb; Table 1, Fig S1 and Table S1). The longest scaffold increased from 58.28Mb to 73.92Mb. Importantly, 539 original draft assembly scaffolds were reduced to just 33 scaffolds, which now contained half of the genome (Fig S2).

**Table 1.** Details of the draft assembly compared to the HiRise assembly.

| Assembly details | Draft | HiRise |
| --- | --- | --- |
| Average coverage | 495.1x | 52.6x |
| Total length (Mb) | 2,478.4 | 2,478.4 |
| **Contigs** |  |  |
| No. of contigs | 153,485 | 153,521 |
| Contig N50 (kb) | 44.137 | 44.131 |
| **Scaffolds** |  |  |
| No. of scaffolds (≥ 0) | 12,483 | 10,007 |
| Longest scaffold (Mb) | 58.28 | 73.93 |
| Scaffold N50 (Mb) | 8.19 | 22.6 |
| No. of scaffolds ≥ N50 | 80 | 33 |
| Scaffold N90 (Mb) | 1.13 | 3.33 |
| **Gaps** |  |  |
| Number of gaps | 141,005 | 143,517 |
| Percent of genome in gaps | 6.75% | 6.76% |

### Identification of genetic variants

We next applied a modified ddRAD sequencing protocol previously described in [24] to generate libraries for 153 13-lined ground squirrels from which we obtained DNA from frozen tissue. After aligning the resulting sample library reads to the HiRise genome assembly (Table S2), we retained 337,695 loci (50.65 Mbp) that fell between predicted *BglII* and *DdeI* target regions, with coverage of at least one read in one individual. Applying variant calling and filtering to these loci (Fig S3), we next identified 786,453 biallelic variants, which had an overall Ti/Tv ratio of 2.19, comparable to ratios reported within intronic and intergenic regions [25]. For use in downstream analyses, we retained 575,178 variants for which genotypes were present in at least 90% of the individuals. Of these retained variants, 35,257 were indels, whereas 539,921 were single nucleotide polymorphisms.

### Population Structure and Genetic Relatedness

The squirrels genotyped in this study originated from wild stock trapped in disparate geographical locales (Fig 1D). The records for their exact source and relatedness were not always available. This was not due to intentional sampling design, but rather due to the availability of squirrels each year, either trapped from the wild or supplied from a breeding colony, and the biological questions originally being pursued. Therefore, to identify population structure within our sample set, we applied ADMIXTURE clustering with 5-fold cross validation [26] on *K*=2 through *K*=10 ancestral populations using a set of 54 unrelated individuals who best represented the ancestries of all squirrels (Fig S4; see Methods). We then applied ADMIXTURE projection to estimate proportions of learned ancestries within the remaining 99 squirrels. The lowest cross validation error occurred at *K*=3 (Fig 2A, top plot), where individuals separated into Colorado (CO), Illinois (IL) and Wisconsin (WI) components. The pairwise genetic distance (F_ST_) estimates between populations were 0.47 and 0.31 for CO vs. WI and IL, respectively, and 0.30 for WI vs. IL, indicating moderate to strong genetic drift. The individual home range for a 13-lined ground squirrel is 0.01 – 0.05 km^2^ [27]. Observed genetic differences may simply be due to isolation by geographic distance [28].

**Fig 2.**
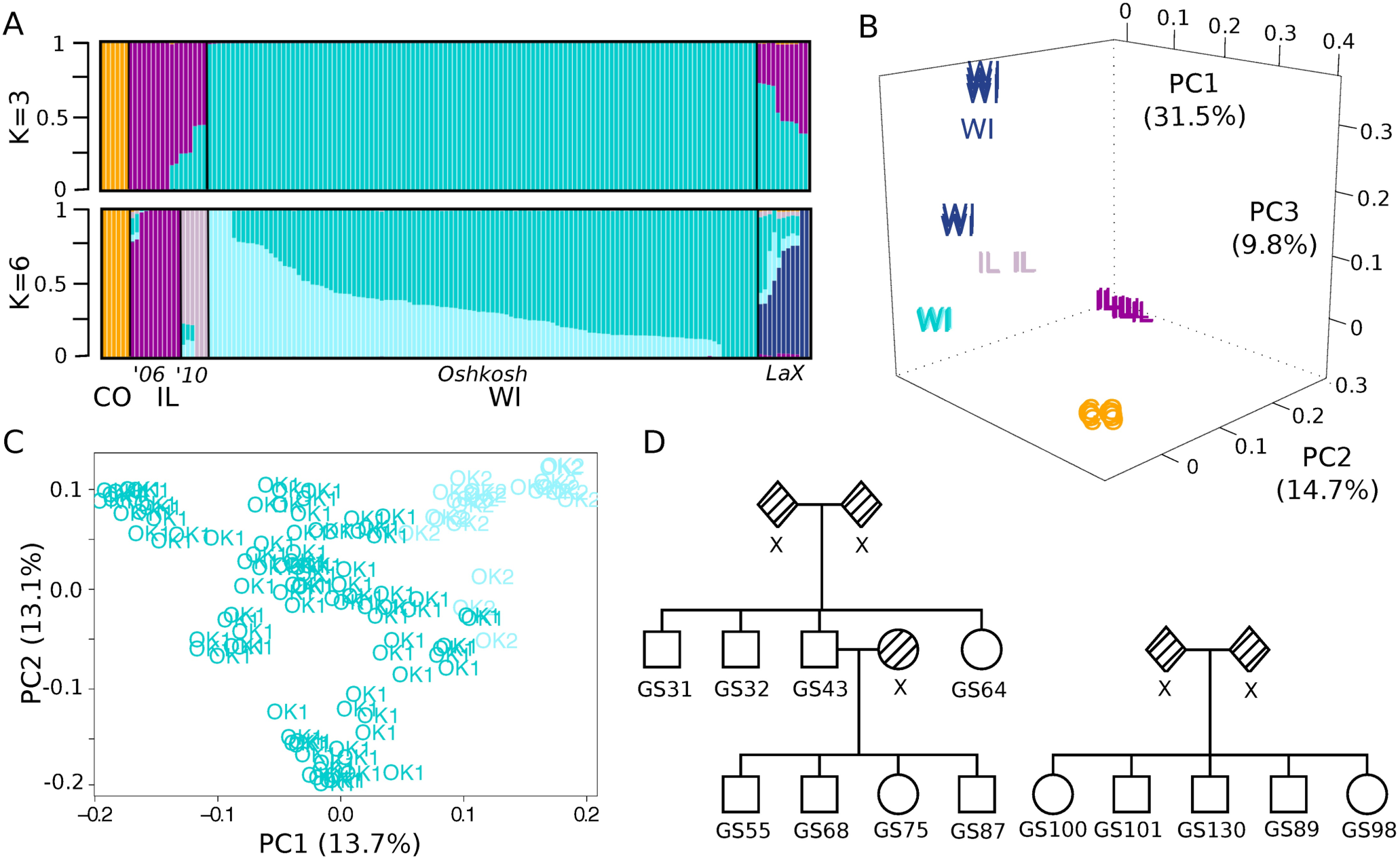
Genotype-by-sequencing reveals population structure and relatedness among sampled squirrels. (A) ADMIXTURE analysis results showing *K*=3 or *K*=6 genome-wide specific ancestry estimates. Squirrels are shown as vertical bars with proportion of specific ancestry colored within each bar. Populations are clustered and labeled by geographic sampling locales (U.S. state, and for WI, city) and for those from IL, sampling years. (B) Principal components analysis of all 153 genotyped squirrels. The first 3 PCs are plotted, with individuals labeled by state and colored by the population for which they have the greatest proportion of ancestry as determined by *K*=6 ADMIXTURE analysis shown in (A). (C) Principal components analysis of 119 squirrels from the Oshkosh WI population. The first 2 PCs are plotted with individuals labeled and colored by Oshkosh sub-population (*OK1* or *OK2*) as determined by *K*=6 ADMIXTURE analysis shown in (A). (D) Representative pedigrees reconstructed from identity by descent (IBD) and kinship coefficient estimates of the Oshkosh squirrels. Shaded shapes labeled “x” indicate relatives not genotyped in this study.

At *K*=6, we observed separation most consistent with records about sampling (Fig 2A, bottom plot). For instance, the algorithm identified a La Crosse, WI (LaX) ancestral component for the squirrels supplied from the UW Oshkosh breeding colony in 2010, matching the breeding records for that year. Additionally, the algorithm identified two ancestral components for the IL squirrels: those purchased in 2006 (’06) belonged to a single ancestry, while those from 2010 (’10) segregated into another ancestry, suggesting different trapping locales between years. While records about the origins of the UW Oshkosh squirrels supplied prior to 2010 were unavailable, the algorithm identified two ancestral components for this breeding colony. The pairwise F_ST_ values were still consistently high (0.33-0.48) among all populations (Table S3), except for the two (non-LaX) within Oshkosh (0.16) and the two within IL (0.23), again supporting the notion of limited gene flow at increased geographical distance.

The first three principal components (PCs) from a PCA-based analysis recapitulated both the observed ADMIXTURE *K*=6 clustering and the known geographical sampling locales of the squirrels (Fig 2B). All populations were distinctly separated, except for the two within Oshkosh, whose separation was only observed at the higher PCs (PC17 and PC19, Fig S5).

### Genetic relatedness within the Oshkosh breeding colony

Due to the strong population structure, and hence large differences in allele frequencies, we limited further analysis to just the Oshkosh population of squirrels (not including LaX, *n*=119), for which we were able to collect the most phenotypic measurements (*n*=72, Table S4) from analysis of the body temperature telemetry data, as opposed to fewer than *n*=10 phenotypic measurements in each of the remaining populations. Our existing records from the breeding colony suggested that many of these squirrels were littermates, although exact relatedness was unknown. We therefore estimated relatedness, adjusting for population substructure with the first PC [29,30], which distinguished the two ancestral components (Fig 2C). Using pedigree reconstruction [31], we identified 19 first-degree families, to which 80% of the squirrels belonged. Consistent with our records, most of these families were composed solely of littermates (Fig 2D, right plot), although in some cases we also identified parent-offspring relationships (Fig 2D, left plot).

### Heritability Estimates for Timing of Autumn Torpor Immersion

We next investigated the effect of genetic architecture on autumn torpor onset within the Oshkosh subset of animals. We estimated heritability of this trait using a linear mixed model, in which we controlled for sex, year of monitoring and date of placement into the hibernaculum (our fixed effects, see methods), and we input the genetic relatedness estimates as the random effect. Unexpectedly, this model converged with no residual error in the variance components, resulting in an estimate of 100% heritability (LMM, Table 2). To confirm this high estimate, we fit a separate Bayesian multivariate general linearized mixed model with the same fixed and random effects. Here, the posterior mode of heritability was 99%, and the confidence intervals were between 61% and 99.9% (MCMCgrm, Table 2); even the lower bound of the estimate still indicated high heritability, confirming our hypothesis that underlying genetic architecture drives variation in the onset of torpor.

**Table 2.**
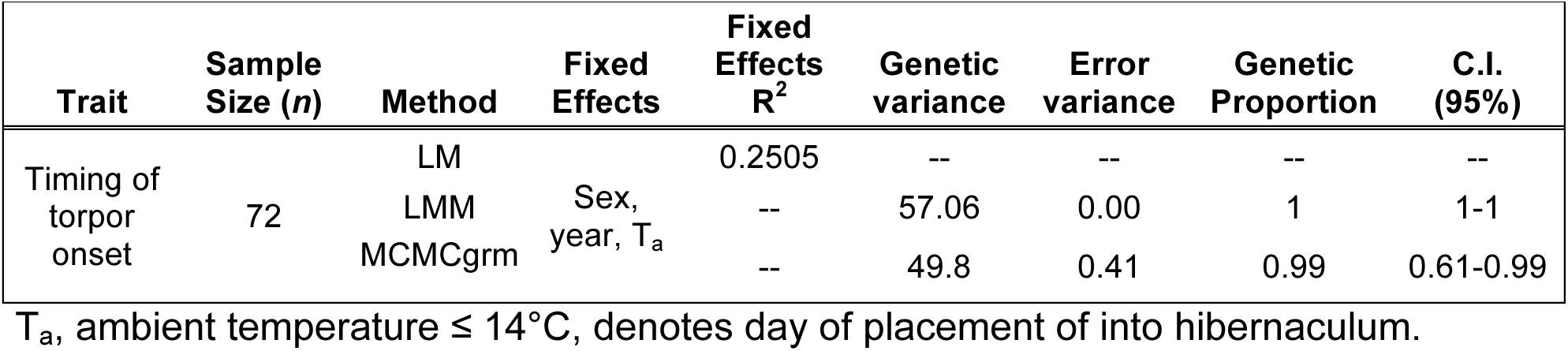
Estimates of heritability for the timing of the first of torpor bout in autumn.

| Trait | Sample Size (n) | Method | Fixed Effects | Fixed Effects R<sup>2</sup> | Genetic variance | Error variance | Genetic Proportion | C.I. (95%) |
| --- | --- | --- | --- | --- | --- | --- | --- | --- |
| Timing of torpor onset | 72 | LM |  | 0.2505 | -- | -- | -- | -- |
|  |  | LMM | Sex, year, T<sub>a</sub> | -- | 57.06 | 0.00 | 1 | 1-1 |
|  |  | MCMCgrm |  | -- | 49.8 | 0.41 | 0.99 | 0.61-0.99 |

T<sub>a</sub>, ambient temperature ≤ 14°C, denotes day of placement of into hibernaculum.

### Genome-wide Association Scan

We identified genetic variants associated with the onset of autumn torpor by performing a genome-wide association scan (GWAS) using 46,996 variants with a minor allele frequency (MAF) ≥ 0.05 and the fit from the linear mixed model. As this was an exploratory analysis using a relatively small sample set, we set a significance cut-off at p≤5 ×10^−4^. After accounting for LD, we identified 21 loci that we considered significantly associated with the phenotype (Fig 3A and Table 3). Although none of the variants met strict genome-wide significance after Bonferroni correction (p<1×10^−6^), a plot ofthe observed vs expected quantiles of log-transformed p-values (Q-Q plot) showed an excess of significant values well above the dashed line in the tail of the distribution (Fig 3B). Furthermore, while the estimated mean allelic effect size was −0.04 days (SD=1.84, *n*=46,996), the effect sizes for these 21 significant variants were all within either the top or bottom 1 % of the total distribution, being at least ± 4.25 days for each additional allele (Fig 3C and Table 3).

**Fig 3.**
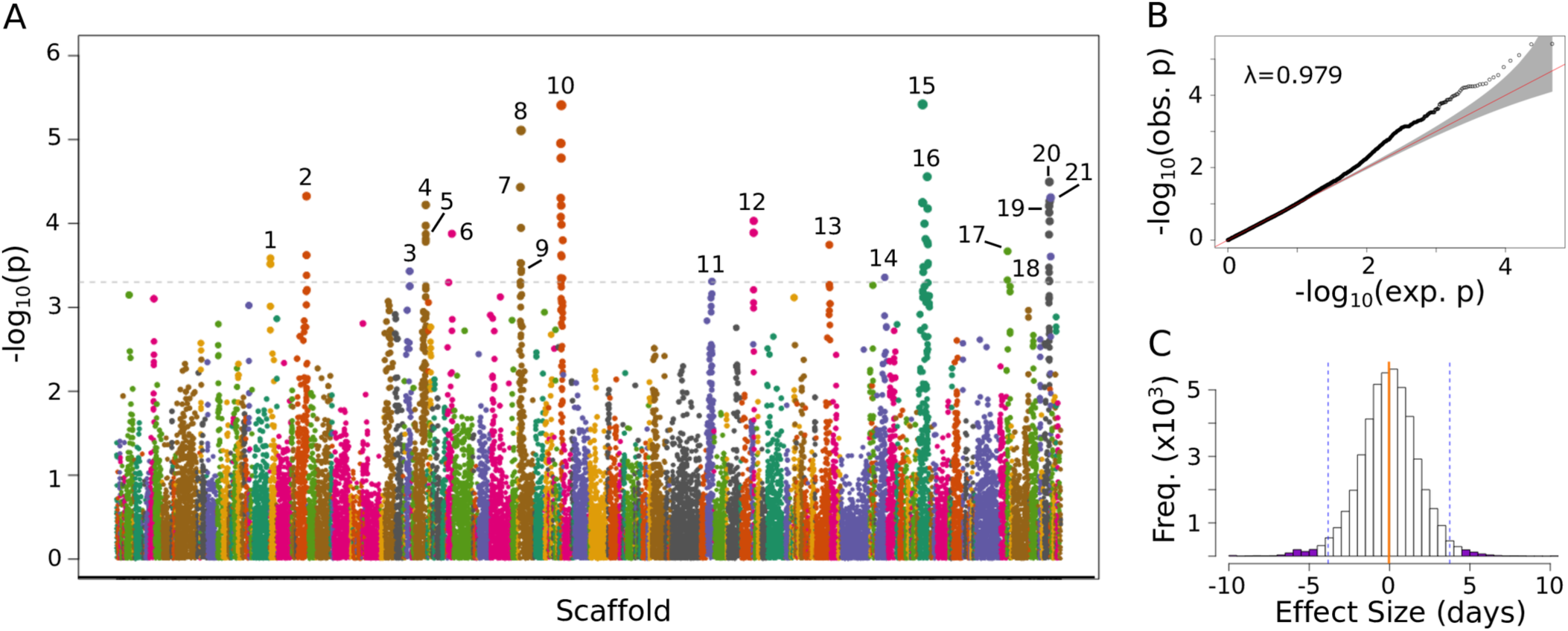
GWAS identifies genetic variants significantly associated with date of first torpor in 13-lined ground squirrels. (A) Manhattan plot shows the negative log-transformed p-values of 46,996 variants (MAF>0.05) tested for association with date of first torpor in 72 squirrels. Variants are ordered by position on scaffold, which are colored along x-axis. Dashed line indicates cutoff for significance (p<5×10^−4^). Significantly associated variants, pruned for LD, are numbered and correspond to those detailed in Table 3 and in Figs 5 and 6. (B) Q-Q plot of the GWAS log-transformed p-values. (C) Histogram of effect sizes of the 46,996 variants on date of first torpor. Orange vertical line marks the mean, dashed vertical lines mark upper and lower bounds of 98^th^ percentile and purple shading indicates effect sizes of the 21 significantly associated variants.

**Table 3.**
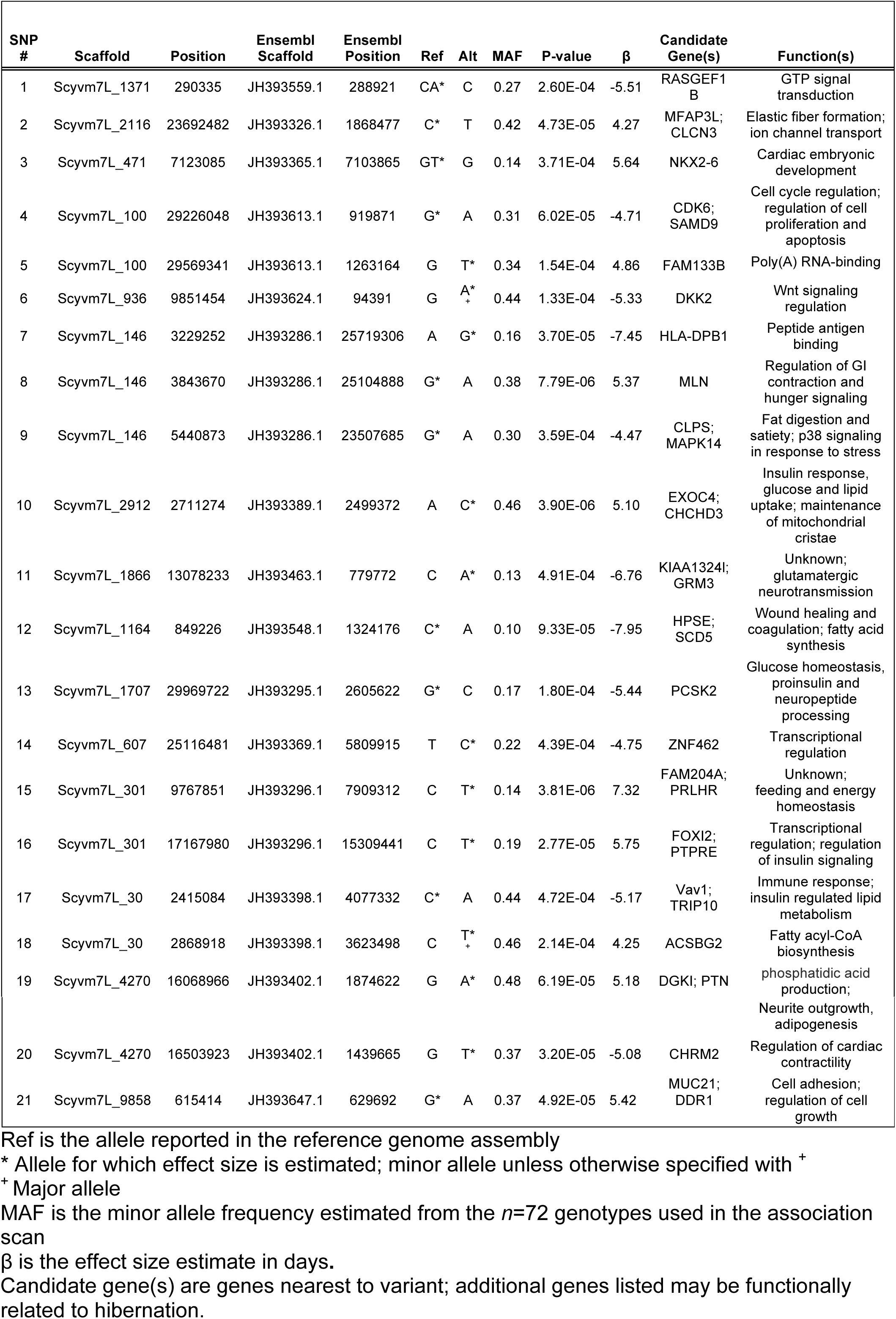
Details about the GWAS variants significantly associated with the onset of torpor

We estimated the amount of phenotypic variance explained by the significant loci via linear regression. While the initial model fit with just the fixed effects of sex, year of monitoring and date of hibernaculum placement accounted for 25% of the variance in onset of torpor (Fig 4A), these 21 markers explained 54% of the variance (Fig 4B) and when combined with the fixed effects, accounted for 85% of the total variance in the phenotype (Fig 4C). Furthermore, the most significant GWAS variant (SNP 15, Table 3) alone accounted for 21% of phenotypic variance, while the subset of the top five most significant loci (SNPs 15, 10, 8, 16 and 20) explained 47.5% of the variance, excluding fixed effects. Hence, a small subset of markers accounted for most of the genetic component underlying the timing of autumn torpor in this population of 13-lined ground squirrels.

**Fig 4.**
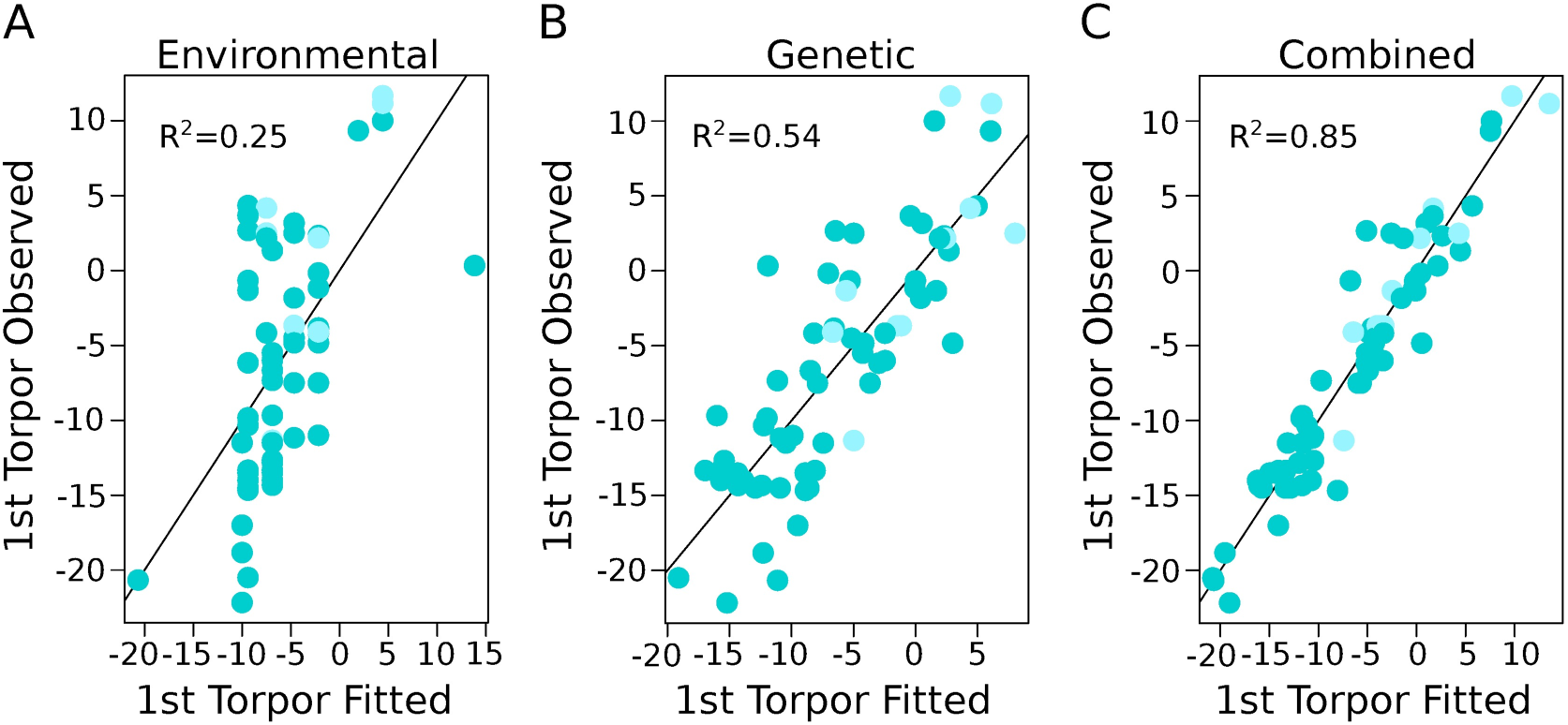
Combined environmental and genetic effects account for variation in the start of torpor. Plots show correlation between fitted and observed values for the start of torpor (in days from hibernaculum placement) using a linear regression model. Adjusted R^2^-value is labeled in each. Shading matches Figs 2A and 2C. (A) Linear model fit with the environmental variables of year of monitoring and date of hibernaculum placement, and the biological variable of sex (i.e. the “fixed effects”, see methods). (B) Linear model fit with the 21 significant SNP genotype combinations for each squirrel. (C) Linear model fit with the variables from both (A) and (B).

When we examined the genes located nearest these significant variants, many were functionally related to themes consistent with physiology underlying the transition to hibernation, such as insulin processing and signaling, feeding and satiety, and control of heart rate (Table 3). Intriguingly, the most significant variant, SNP 15, was located nearest the gene *family with sequence similarity 204 member A* (*FAM204A*), whose function is poorly characterized (Fig 5A). However, the *prolactin*-*releasing hormone receptor* (*PRLHR*), involved in stimulating prolactin release and control of feeding [32], and hence more consistent with roles in circannual timing and hibernation, was approximately 270kb from this marker. The second-most significant marker, SNP 10, was located between two genes that are also functionally relevant within the scope of hibernation: *coiled-coil-helix-coiled-coil-helix domain containing 3* (*CHCHD3*) and *exocyst complex component 4* (*EXOC4*; Fig 5B). While *CHCHD3* maintains the structural integrity of mitochondrial cristae [33], *EXOC4* is a component of the exocyst complex involved in the secretion of insulin [34], as well as lipid and glucose uptake in response to insulin signaling [35,36]. Two weakly-linked variants (*r*^2^=0.33) located approximately 500kb from each other, SNP 19 and SNP 20 (Figs 5C & 5D), were near *pleiotrophin* (*PTN*), a growth factor involved in neurogenesis and axonal outgrowth, angiogenesis and adipogenesis [37-39], and the *muscarinic acetylcholine receptor M2* (*CHRM2*), which mediates bradycardia in response to parasympathetic-induced acetylcholine release [40], a phenomenon well characterized in hibernation [41,42]. Finally, *motilin* (*MLN*), a small peptide hormone that regulates gastrointestinal contractions and stimulates hunger signaling [43] was nearest SNP 8 (Fig 5E), while *prohormone convertase 2* (*PCSK2*), an enzyme that activates hormones and neuropeptides [44-46], including cleavage of proinsulin into its mature form [47], was located in close proximity to SNP 13 (Fig 5F).

**Fig 5.**
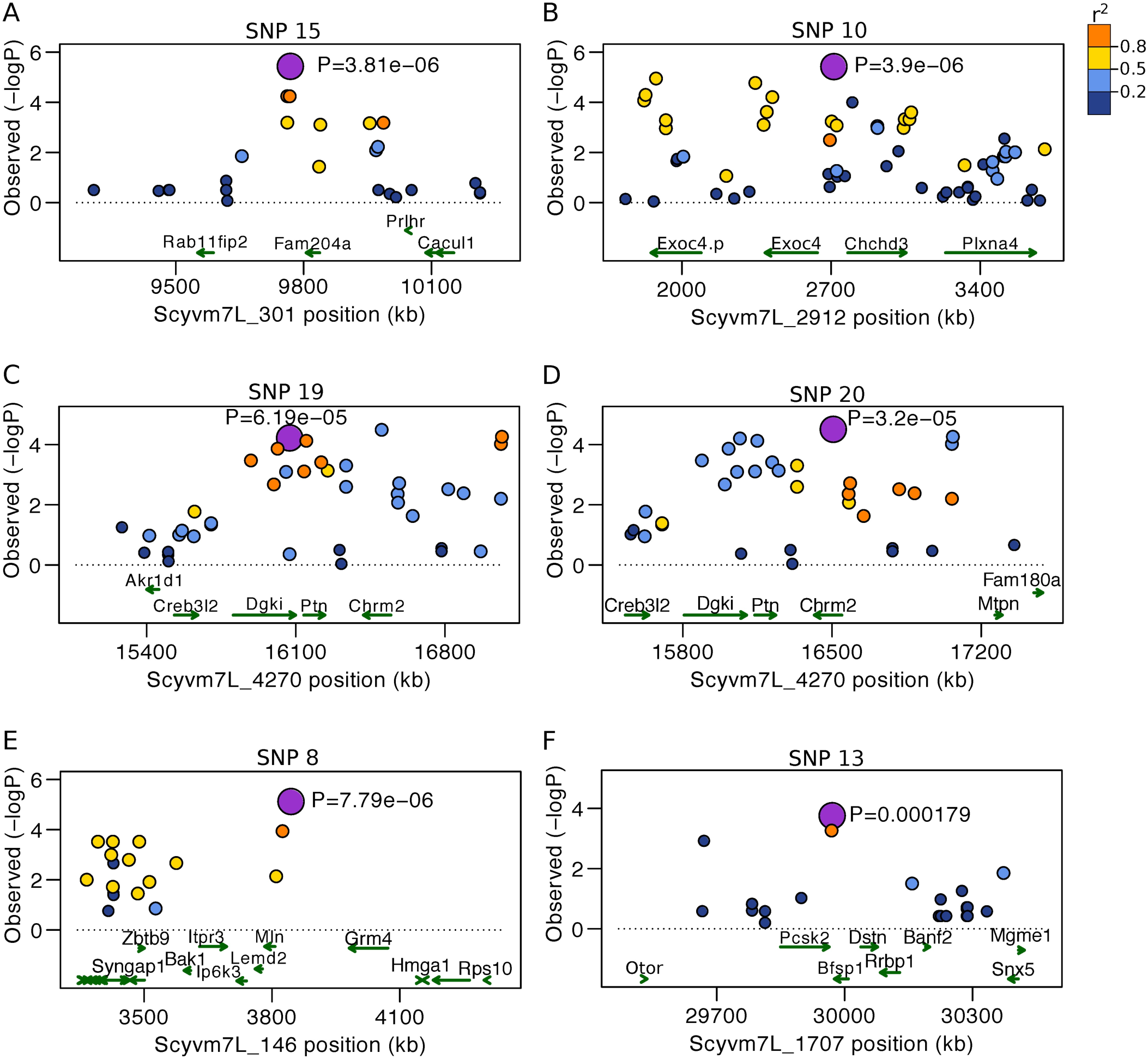
Regional Manhattan plots show locations of selected significant SNPs in proximity to nearest genes. Each plot is centered on one of the significant SNPs, which are labeled on top by number (see Fig 3A and Table 3) and shaded purple. Other variants within region are colored by LD value (*r*^2^) in relation to the significant SNP. Genes are shown below as green arrows and labeled by gene symbol. (A) SNP 15, the most significant SNP in the GWAS, is nearest *FAM204A*. (B) SNP 10, the second most significant SNP, is between *CHCHD3* and *EXOC4*. (C) SNP 19 is located nearest *DGKI* and *PTN*. (D) SNP 20 is located nearest *CHRM2.* (E) SNP 9 is nearest *MLN*. (F) SNP 13 is between *PCSK2* and *BFSP1*.

### Identification of eQTLs using transcriptomic datasets

We hypothesized that these significant loci might be linked to gene regulatory variants. We therefore applied an expression quantitative loci (eQTL) analysis using the EDGE-tag transcript datasets from heart, liver, skeletal muscle (SkM) and brown adipose tissue (BAT) [48,49], since a subset of the squirrels genotyped in this study were assayed for transcriptome expression in these prior studies. Under an additive linear model, we identified significant *cis-* eQTL associations (±500kb, q<0.1) for 9/21 variants (Table 4). The most significant GWAS variant, SNP 15, was also the most significant *cis*-eQTL (q<4.4×10^−8^), where the minor allele, associated with a later onset of torpor (Fig 6A, left plot), was correlated with increased expression of *FAM204A* in BAT (Fig 6A, middle-left and middle-right plots) and SkM. Several variants were associated with expression changes in the previously identified candidate genes (Table 3). SNP 19, associated with a later torpor onset (Fig 6B, left plot), was also correlated with decreased expression of *PTN* in BAT (Fig 6B, middle-left and middle-right plots), while SNP 20, associated with an earlier torpor onset (Fig 6C, left plot), was correlated with increased *CHRM2* expression in heart (Fig 6C, middle-left and middle-right plots). Two variants were correlated with changes in expression of the same transcript in the same direction across tissues: in heart and SkM, SNP 5 correlated with increased expression of *family with sequence similarity 133 member b* (*FAM133B*), while SNP 21 correlated with decreased expression of *coiled-coil alpha-helical rod protein 1* (*CCHCR1*; Table 4). In contrast, several variants were associated with expression changes of different transcripts depending on the tissue examined, suggesting their linkage to several regulatory loci within the region or to a shared regulatory site that exerts its effects on multiple nearby genes [50]. For example, SNP 2 was associated with decreased *chloride voltage-gated channel 3* (*CLCN3*) expression in SkM, yet also associated with increased *microfibril associated protein 3 like* (*MFAP3L*) expression in heart (Table 4).

**Fig 6.**
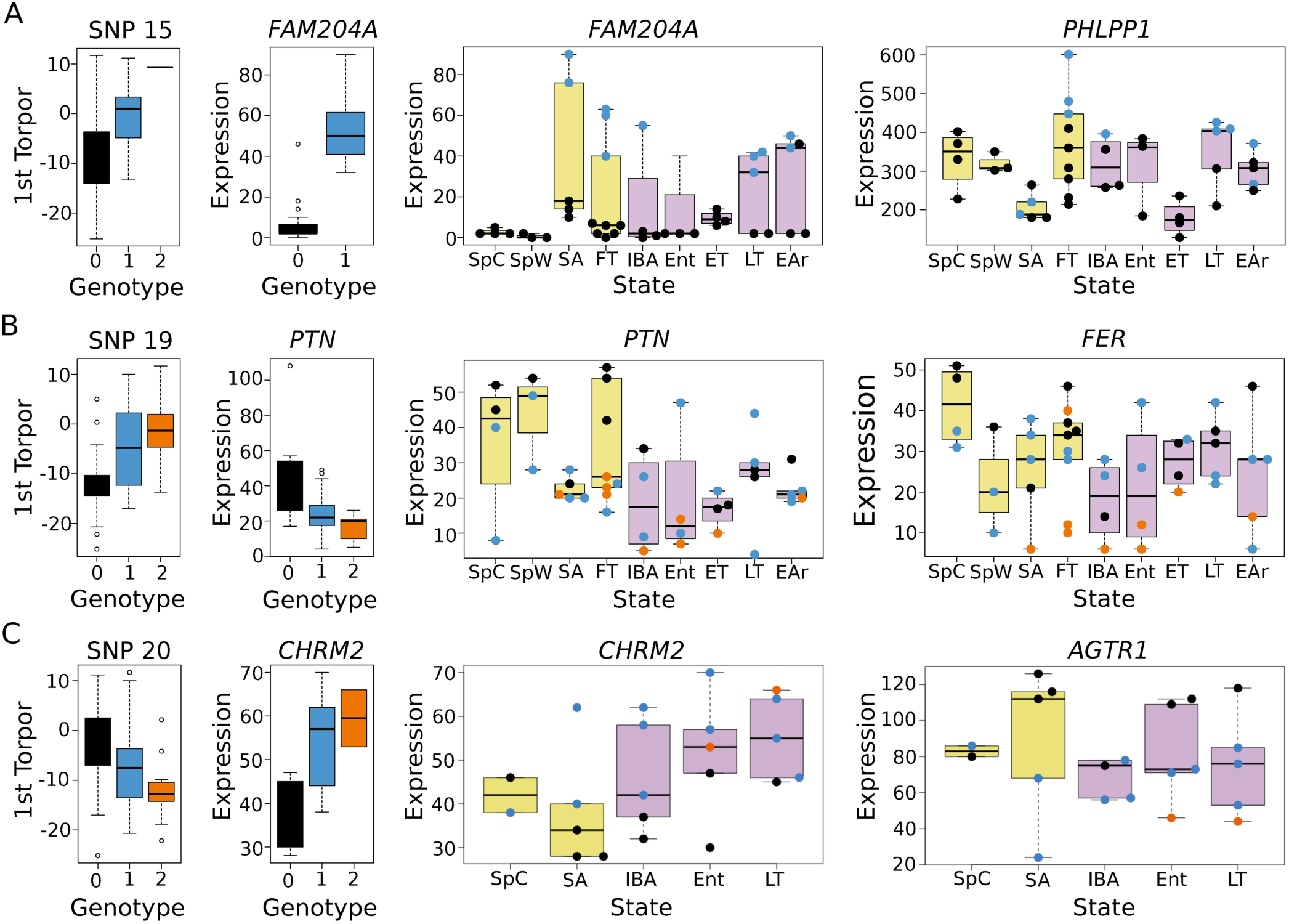
Significant GWAS SNPs are also *cis-* and *trans-*eQTLs that explain variation in mRNA expression. (A) Effect of SNP 15 genotype on date of first torpor (far left) and expression of its c/s-eGene, *FAM204A*, in BAT (middle left). Middle right plot shows effect of genotype on transcript expression within each of the nine distinct physiological and seasonal states interrogated in the original transcriptome study (labeled below, see methods for explanation of sampling abbreviations). Shaded yellow boxes indicate physiological states from homeothermic and transitional portions of hibernator’s year (spring-autumn), while purple shaded boxes are those from within deep hibernation. Far right plot shows effect of genotype on expression of the most significant *trans-eGene*, *PHLPP1*. (B-C) Labeling is as in panel (A).(B) Effect of SNP 19 on *cis*-eGene *PTN* and *trans*-eGene *FER* in BAT. (C) Effect of SNP 20 on *cis*-eGene *CHRM2* and *trans*-eGene *AGTR1* in heart.

**Table 4.**
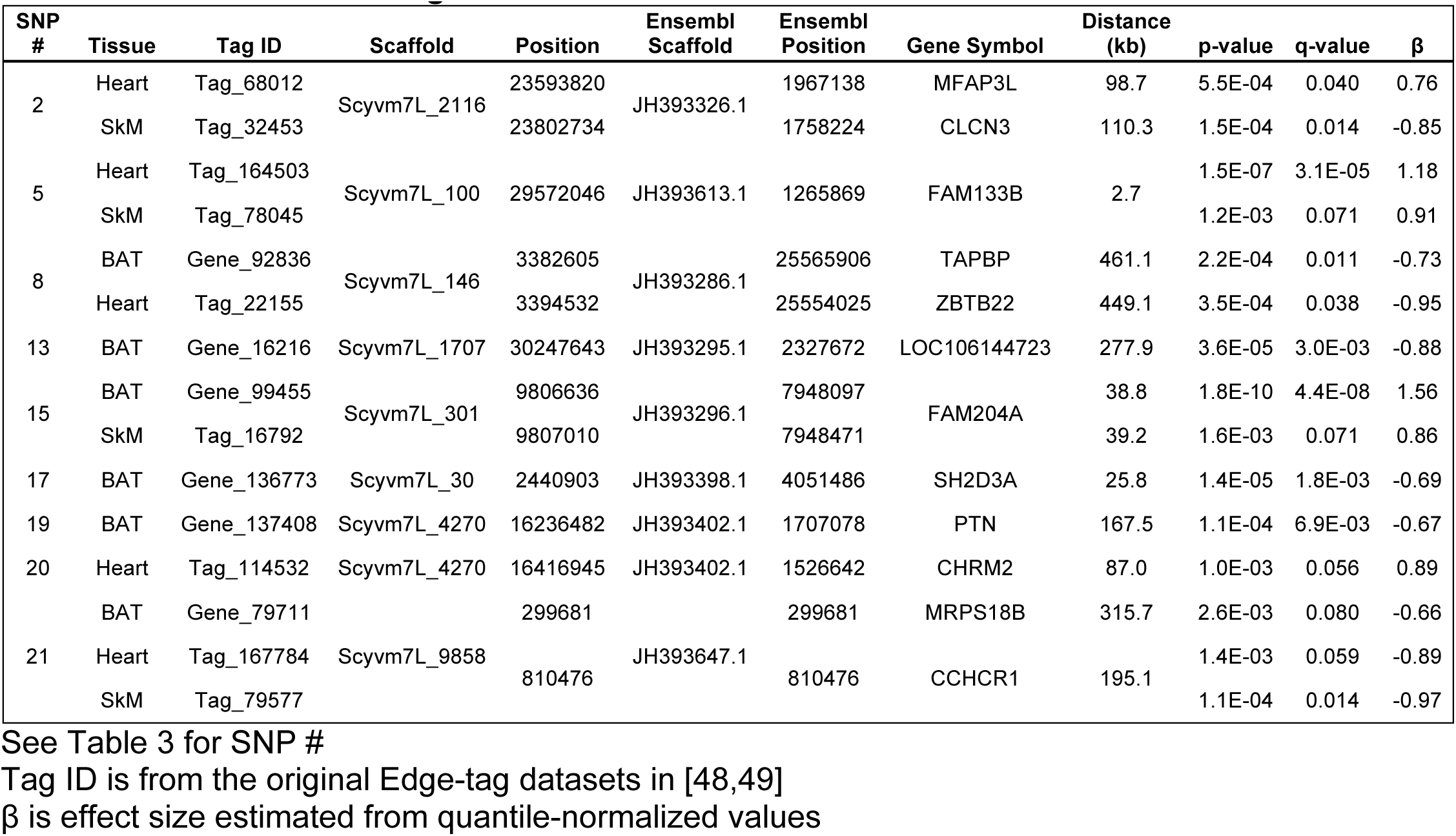
Details about the significant *cis*/-eQTLs

None of the variants met significance thresholds (q<0.1) to be identified as *trans-eQTLs* (Tables S5-S8); however, this was likely due to the relatively small sample sizes (*n*=22-23 in heart, SkM and liver; *n*=43 in BAT) and the large number of EDGE-tags tested (25-30K per tissue) in each dataset. We therefore examined the top *trans*-eGene (>500kb from variant) for each significant cis-eQTL, hypothesizing that we would identify genes within the same pathway or consistent with the physiology of the cis-eGene. Indeed, in heart, the *angiotensin II receptor type 1*, *AGTR1,* involved in regulation of blood pressure [51], was the top *trans-eGene* for SNP 20 (p=2.97×10^−5^, Table S5). In contrast to *CHRM2*, this transcript showed decreased expression in relation to the minor allele (Fig 6C, right plot), consistent with the physiology of torpor, where reduced heart rate is coupled with decreased blood pressure [52,53]. In BAT, the top *trans-*eGene for SNP 19 was *FER Tyrosine Kinase* (*FER*; p=4.73×10^−5^, Table S8), whose expression, like *PTN*, was also decreased in relation to the minor allele of its SNP (Fig 6B, right plot). Both *PTN* and *FER* phosphorylate β-catenin [54,55], suggesting a role for this pathway in the start of torpor. Finally in BAT, the top *trans*-eGene for SNP 15 was *PH domain and leucine rich repeat protein phosphatase 1* (*PHLPP1;* p=2.1×10^−5^, Table S8), whose expression, like *FAM204A*, was increased in relation to the variant (Fig 6A, right plot). *PHLPP1* is a protein phosphatase that dephosphorylates and inactivates both *Akt2* and *protein kinase C* [56]. *Akt2* is expressed highly in insulin-responsive tissues, including BAT, where it modulates glucose uptake and homeostasis [57], as well as non-shivering thermogenesis (NST) [58]. Moreover, increased *PHLPP1* expression is associated with insulin-resistance and hyperinsulinaemia [59]; hence, this gene may play a role in the switch from glucose to fat-based metabolism that occurs at the onset of seasonal heterothermy, and more specifically in BAT, the regulation of NST.

## Discussions

Mammalian hibernation is a highly dynamic and extraordinary phenotype that remains poorly understood. While it has been characterized at behavioral, whole body, cellular, and molecular levels, a genetic basis of the phenotype has yet to be established. Our study is the first, to our knowledge, to characterize genome-wide variation within a hibernator, the 13-lined ground squirrel. This enabled us to estimate the heritability of, and identify genetic variants associated with, the onset of seasonal heterothermy.

Our results of heritability are consistent with those from a study that reported significant heritability in spring emersion from hibernation in wild Columbian ground squirrels [60]. However, our estimates for immersion into hibernation are much higher. This is likely due to differences between monitoring animals in an animal facility, where environmental conditions and access to food are relatively constant and social cues are minimal, and monitoring animals in the field, where phenotypic plasticity in response to changing environmental conditions also influences phenological timing [61,62]. In the wild, differences in age (adult vs juvenile) and hibernation timing have been observed [61]. While age did not significantly affect hibernation onset in our study, it is worth noting that 74% of the squirrels in this dataset were juveniles, and therefore hibernation-naïve prior to their first torpor bout. Juveniles likely face a far greater challenge in growing and fattening sufficiently to support winter hibernation in the wild; in contrast, in a relatively constant, resource-rich laboratory environment, hibernation onset for these animals may be particularly driven by endogenous mechanisms, thus increasing heritability estimates.

In contrast to complex human diseases, where often thousands of variants of relatively small effect influence complex physiological phenotype [63], we find that relatively few loci of large effect account for phenotypic differences in the seasonal onset of hibernation. Our results are comparable to what has been observed for adaptive traits in *A. thaliana* [64], morphological variation in domesticated dogs [65] and plate armor of threespine sticklebacks [66]. This may be simply due to limited sample size, as we were underpowered to detect variants with small effect size and/or at low frequency. However, the results from our ADMIXTURE analysis suggest that wild-trapped founders from two distinct populations were crossed to form the breeding colony. One possible explanation is that torpor onset differs between these two populations, and if so, local genetic adaptation in each could explain the relatively few loci of large effect driving phenotypic variation [67].

Although more research is now needed to determine the precise role in torpor immergence for each of these loci, we propose candidate genes, oftentimes the closest to the marker, due to their function being closely related to the physiology of this seasonal transition. In particular, several genes are known to modulate food intake, such as the *prolactin-releasing hormone receptor* (*PRLHR*), *motilin* (*MLN*), and *procolipase* (*CLPS*) [68]. Researchers have hypothesized that mechanisms governing food intake and metabolic suppression are linked, and that hibernation cannot begin until food intake has ceased [7,69]. Our results support this hypothesis, and present new candidates for study in hibernation.

Perhaps the most intriguing candidate gene is *PRLHR.* Sharing a common ancestry with the *NPY* receptors [70], this receptor is expressed primarily in the anterior pituitary, as well as in distinct regions of the brain, including the hypothalamus. Its knockout in mice results in an obese, hyperphagic phenotype, while administration of its agonist, prolactin-releasing hormone (*PRLH*), induces hypophagia and decreases body mass [32]. Both *PRLH* and *PRLHR* appear to mediate the effects of leptin, including activation of NST in BAT [71]. Further, *PRLH* belongs to a class of neuropeptides containing a C-terminal RFamide motif [72]; other RFamide-related peptides are involved in circannual regulation of reproduction [73,74]. Binding of *PRLH* to *PRLHR* may induce prolactin release, a hormone that has established roles in the timing of circannual rhythms, such as seasonal molt [75] and reproduction [76]. Moreover, serum prolactin levels coincide with resumption of post-hibernation feeding in marmots [77]. Thus, we hypothesize that *PRLHR* is involved in the regulation of circannual food intake in hibernators; its role in this rhythm warrants further investigation.

However, we note that the most significant *cis*-eGene for SNP 15 was not *PRLHR*, but rather its near neighbor, *FAM204A*. This may be due to the marker being linked to several regulatory sites within the region or to a single regulatory site that affects multiple nearby genes. Additionally, *PRLHR*, along with other candidate genes, was not expressed in the tissues for which we had transcriptome data, a limiting factor in our analysis. A role for *FAM204A* in hibernation is unclear, as not much is known about this gene. It appears to be expressed in every tissue and localizes to the nucleus (www.proteinatlas.org [78]), where it interacts with a histone acetyltransferase and a methyltransferase [79]. Therefore, it may play a role in epigenetic regulation of gene expression, possibly in response to the onset of fasting and a switch to fatty acid metabolism [80], as animals prepare for hibernation in the fall.

Finally, our results highlight the power of integrating genome data with transcriptome and other high-throughput data to better understand the genetic mechanisms underlying hibernation. Prior “omics” screens (e.g. [10,12,13,22,48,49,81]) have identified hundreds to thousands of genes differentially expressed among the seasonal and physiological states of the hibernator’s year. While leading to insight into the pathways involved, the results of these screens do not distinguish between genes driving vs. those responding to changes in phenotype. They are also limited to the tissue and time-points being examined and may therefore miss important regulators of phenotype. It is worth noting that while several *cis*-and *trans*-eGenes identified here have clear roles in the physiology of torpor, such as *CHRM2* and *AGTR1*, neither of these were identified in their original studies that screened only for differential expression. Thus, by applying complementary genetic mapping approaches, current limitations inherent to gene expression screening strategies will be addressed and enable new insight into the mechanisms driving hibernation. The approaches used here can be extended to a wide variety of hibernators and the quantifiable components that comprise this highly dynamic phenotype.

## Materials and Methods

### Animals

Animals were procured and housed at the University of Colorado, Anschutz Medical Campus, as previously described [15]. All animal use was approved by the University of Colorado, Anschutz Medical Campus, Animal Care and Use Committee.

Briefly, 130 colony-bred animals (68 females and 62 males) were obtained from the University of Wisconsin, Oshkosh [82] in the summers of 2007—2010 (Fig 1D, “WI”). These included 73 juveniles naïve to hibernation in the year of study, and 57 adults with at least one year of hibernation. While most from the colony were bred from squirrels originally wild-trapped in northeastern Wisconsin (in and around Oshkosh), several of those obtained from the Oshkosh colony in 2010 were actually bred from either a single or both parents wild-trapped in far western Wisconsin, more than 100 miles away (in and around La Crosse, WI). However, records to identify these specific squirrels were not always maintained. In addition, 17 squirrels (9 females and 8 males; ages unknown), wild-trapped in different locales around central Illinois, were obtained from a commercial supplier (TLS Research, Bloomington, IL) in the summers of 2006 and 2010 (Fig 1D, “IL”). Finally, 6 squirrels (3 females, 3 males; ages unknown) were wild-trapped in the summers of 2006 and 2009 in Elbert County and Larimer County, Colorado (5 and 1, respectively; Fig 1D, “CO”).

Upon arrival, animals were housed individually in rodent cages (Fig 1B, bottom) under standard laboratory conditions (20±2°C and 14:10 light-dark cycle, fed cat chow supplemented with sunflower seeds *ad libitum*). In late August or early September, animals not yet euthanized for tissue collection were surgically implanted with an intraperitoneal datalogger (iButton, Embedded Data Systems) and/or a radiotelemeter (VM-FH disks; Mini Mitter,Sunriver, OR) for remote body temperature (T_b_) monitoring until tissue collection. The dataloggers recorded T_b_±0.5°C every 20, 30 or 60 min, while the radiotelemeters transmitted T_b_±0.5°C every 20 sec.

In late September or early October, the squirrels were moved to the hibernaculum to facilitate hibernation. The temperature was lowered stepwise over a two-week period to 4°C. Food was removed as animals became torpid.

### Tissue Collection and Telemeter Retrieval

Liver samples were collected at different points throughout the year for use in other biochemical studies as previously described [19,81,83,84]. All animals were exsanguinated under isoflurane anesthesia, perfused with ice-cold saline, decapitated, and dissected on ice; tissues were immediately snap frozen in liquid nitrogen and stored at -80°C until processed further. Telemeters were retrieved during tissue collection.

### Body Temperature Telemetry Analysis

To identify the first day of torpor, the telemetry data were analyzed in R [85]. T_b_ was averaged over 4-hour windows. Homeothermic T_b_ typically ranged from 34-39°C. The first torpor bout was defined as the first point at which T_b_ fell to or below 25°C (approximately 3-5°C above ambient prior to hibernaculum placement, Fig 1C). Most telemetry data continuously logged T_b_ from the beginning of September of each year. However, in several cases, telemetry recordings did not start until mid-September. Of these, only cases in which first torpor occurred after a minimum of 10 days of continuous monitoring were included for further analysis. In order to merge data across years, date of first torpor was transformed into date from placement into the hibernaculum, which was centered as day 0; hence, all days prior are negative in value, while post-placement dates are positive.

### HiRise Genome Assembly and Annotation

A Chicago library was prepared as described previously [23] from a single 100mg frozen liver sample. Briefly, ~500ng of high molecular weight gDNA (mean fragment length = >50kbp) was reconstituted into chromatin in vitro and fixed with formaldehyde. Fixed chromatin was digested with *DpnII*, the 5’ overhangs filled in with biotinylated nucleotides, and then free blunt ends were ligated. After ligation, crosslinks were reversed and the DNA purified from protein. Purified DNA was treated to remove biotin that was not internal to ligated fragments. The DNA was then sheared to ~350 bp mean fragment size and sequencing libraries were generated using NEBNext Ultra enzymes and Illumina-compatible adapters. Biotin-containing fragments were isolated using streptavidin beads before PCR enrichment of the library. The library was sequenced on an Illumina HiSeq 2500 (rapid run mode) to produce 150 million 2x101bp paired end reads, which provided 52.6x physical coverage of the genome (1-50kb pairs).

The 13-lined ground squirrel draft assembly, shotgun reads, and Chicago library reads were used as input data for HiRise, a software pipeline designed specifically for using proximity ligation data to scaffold genome assemblies [23]. Shotgun and Chicago library sequences were aligned to the draft input assembly using a modified SNAP read mapper (http://snap.cs.berkeley.edu). The separations of Chicago read pairs mapped within draft scaffolds were analyzed by HiRise to produce a likelihood model for genomic distance between read pairs, and the model was used to identify and break putative misjoins, to score prospective joins, and make joins above a threshold. After scaffolding, shotgun sequences were used to close gaps between contigs. Table S1 describes the input draft assembly scaffold placement within the HiRise scaffolds.

Gene annotations from the Ensembl (Release 86) and NCBI (Release 101) datasets were lifted over to the HiRise assembly using a custom Python script and Table S1.

### Genotype-by-sequencing

We used the modified ddRAD sequencing protocol previously described in [24]. Briefly, high molecular weight DNA was extracted from 8-15 mg of frozen liver with commercially available kits. Digestion and ligation reactions were performed using 200ng of genomic DNA from each sample with *BglII* and *DdeI* and 11-fold excess of sequencing adaptors. Samples were amplified by PCR for 8-12 cycles with a combination of index-containing primers.Between 50-60 samples were pooled in equal amounts according to their concentration of PCR product between 280-480 bp as measured by Tapestation (Agilent Technologies, Santa Clara, CA). Inserts were size-selected on a BluePippin (Sage Science, Beverly, MA) with a target range of 380±100bp and sequenced on the Illumina NextSeq in single-end 151 bp mode using a high output kit.

### Variant calling and Filtering

Reads were mapped to the 13-lined ground squirrel HiRise assembly with BWA v.0. 7.12. [86]. Tables of predicted *BglII* and *DdeI* restriction digest fragments were generated as described in [24], and sequencing coverage was measured at these sites. We then defined “target regions” for variant calling using the set of fragments between 125-350 bp long that had non-zero coverage in at least one individual. The mapping data are summarized in Table S2.

Because publicly available data on 13-lined ground squirrel genetic variation is nonexistent, we instead used several variant callers to identify genetic variants and to assess concordance of the genotype calls at each site. Variant calling was performed independently with Sentieon [87], Platypus [88] and Samtools [89,90]. In Sentieon, the pipeline algorithms indel realignment, base quality score recalibration, haplotyper and GVCFtyper were implemented with default settings. In Samtools, variants were called jointly using mpileup to first compute genotype likelihoods and then BCFtools to call genotypes with default parameters. Finally, variants were called jointly in Platypus with the following parameters: minFlank=3, badReadsWindow=5, maxVariants=12, and minReads=6. Only biallelic variants that both passed the filter flags and were identified by all three callers were retained (Fig S3). These variants were next intersected with GATK [91] and compared for genotype call concordance across samples [92]. Those that were ≥ 95% concordant were kept. Basic statistics about the callset, including depth, missingness, heterozygosity, Hardy-Weinberg equilibrium, and TiTv ratio were calculated in VCFtools [93]. Variants with excessive coverage (≥ 65X, approx. 4x the mean coverage, Table S2) and heterozygosity (obs./exp. ratio ≥ 1.2) were removed from the callset (Fig S3). Sample libraries with excessive missingness and/or heterozygosity were removed, remade, and resequenced. Variants then were reiteratively called and filtered as described above. Finally, variants present in ≥ 90% of the sample libraries were used for further downstream analyses.

### Population Structure and Genetic Relatedness Estimates

We first inferred relatedness from identity-by-state (IBS) estimates among all genotyped squirrels (*n*=153) using KING software [94]. Due to the expectation that the Colorado squirrels are of a separate subspecies [95], relatedness was calculated independently for this subset. We selected an unrelated subset of 54 squirrels that best represented the ancestries of all squirrels within the dataset using the GENESIS package [96] in R [85]. Variants were pruned for LD in PLINK v. 1.9 [97] using the parameters --indep-pairwise 50 10 0.5, which reduced the dataset to 148,870 variants. We then ran unsupervised ADMIXTURE [26] for K=3 through K=10 with 5-fold cross-validation. To estimate the ancestries of the remaining 99 squirrels, we ran ADMIXTURE’s projection analysis using the population structure learned in the initial unsupervised analysis, here with K=2 through K=8 and 5-fold cross-validation.

We performed principal components analyses (PCA) with PLINK using 90,376 LD pruned variants with MAF > 0.01 for the entire dataset and 30,356 LD pruned variants with MAF > 0.01 for the squirrels within the Oshkosh, WI, population (*n*=119), as identified by ADMIXTURE analysis. We extracted the top 20 principal components in each analysis.

Finally, we calculated genetic relatedness among the 119 Oshkosh WI squirrels using the GENESIS package, adjusting for both population substructure and inbreeding with the first principal component [29]. We used the resulting kinship coefficients and identity-by-descent (IBD) estimates to reconstruct and visualize pedigrees among the 1^st^ degree relatives with PRIMUS [31]. We also constructed a genetic relatedness matrix from the pairwise kinship coefficients.

### Genome-Wide Association Scan and Heritability Estimates

All analyses, unless otherwise stated, were performed in R [85]. To identify environmental and biological factors that affected the date of first torpor, we applied a linearregression using variables available from records about the squirrels. In this initial model:

Date of first torpor = f(sex + year of monitoring + date of datalogger implantation + age (juvenile vs. adult) + date of placement into hibernaculum + weight (as last recorded before placement into hibernaculum).

We then pruned factors using step-wise regression until we identified a minimum set that did not significantly reduce the adjusted R-squared value from the initial model, yet also returned a low AIC value. In this final model:

Date of first torpor = f(sex + year of monitoring + date of placement into hibernaculum).

These were our fixed effects.

We carried out a genome-wide association scan (GWAS) on the date of first torpor using GENESIS [96]. We first fit a linear mixed model using the fixed effects and the genetic relatedness matrix as the random effect. We then performed SNP genotype association tests with 46,996 SNPS (MAF≥0.05) and the fit from the linear mixed model. As this was an exploratory analysis, we considered any variant with p≤5×10^−4^ to be significantly associated with the phenotype. To account for LD, We calculated the *r*^2^ values for significant SNPs within the same scaffold using PLINK [97,98]. We removed those in moderate to high LD (*r*^2^≥0.5), reporting only the most significant variant.

We estimated heritability of the first day of torpor from the variance components of the linear mixed model. In addition, we also estimated heritability of this phenotype using a separate Bayesian mixed model with the MCMCgrm package in R [99]. Here we input the same fixed and random effects (i.e. genetic relatedness matrix) as in the linear mixed model. For the prior, we used an uninformative inverse-gamma distribution (with variance, V, set to 1 and belief parameter, nu, set to 0.002) on the variance components. We ran three chains, each with a total of 1,000,000 iterations, a burnin of 100,000 rounds and a thinning interval of 200 rounds. Here, all variables had Gelman-Rubin statistics of 1.00 – 1.01, with the absolute value of all autocorrelations < 0.1 and effective sample sizes between 3682.8 and 5294.5. We combined the 3 chains in order to estimate the posterior mode and confidence intervals of the variance components.

Finally, we estimated the effects of the significant GWAS variants on the onset of torpor. Specifically, we used linear regression with the phenotype as the dependent variable and a matrix of significant variant genotypes, either with or without the fixed effects, as the explanatory variables. We also performed forward stepwise regression using genotype combinations from the top 10 significant variants.

### EQTL analysis

We applied an eQTL analysis to identify transcripts whose expression levels were significantly affected by the GWAS variants. Here, we used the EDGE-tag datasets from heart, liver, skeletal muscle (SkM) and brown adipose tissue (BAT) previously described in [48,49], where total RNA was digested with *NlaIII*, resulting in the generation of ≈27-nt “EDGE-tags” [100], which mapped to the 3’UTR’s of transcripts. The squirrels assayed in these prior transcriptome studies were also genotyped in this study: heart (n=22), liver (n=23), SkM (n=22) and BAT (n=43). Due to small sample sizes, we limited our eQTL association tests to significant GWAS SNPS with MAF ≥ 0.2 in heart, liver and SkM and ≥ 0.1 in BAT, which ensured that a minimum of 9 samples contained at least one minor allele.

Tests for both *cis*- (±500kb) and *trans*-eQTLs were performed with Matrix eQTL [101] under an additive linear model. As the purpose of the original EDGE-tag studies was to identify differentially expressed transcripts among distinct physiological states within the hibernator’s year, we included physiological state as a covariate (5 states in heart, liver and SkM: spring cold, SpC; summer active, SA; interbout-aroused in hibernation, IBA; entering torpor in hibernation, Enf; and late torpor in hibernation, LT; 9 states in BAT, in addition to the those previously mentioned: spring warm, SW; fall transiton, FT; early torpor in hibernation, ET; early in arousal from torpor in hibernation, EAr). In BAT, sequencing platform was also included due to the count bias observed in [49]. Finally, to control for outliers and following recommendations by Matrix eQTL, the counts for each Edge-tag were ranked and quantile-normalized before testing.

## Acknowledgements

We thank members of the Bustamante Lab for their helpful discussion while this research was being conducted. We also thank R. Russell, A. Hindle and members of the Martin lab who assisted with the 13-lined ground squirrel care, surgical implantation of dataloggers and collection of tissue.

C. D. Bustamante is a Chan Zuckerberg Biohub investigator.

## Supporting Information Captions

**Fig S1. Comparison of the contiguity of the input assembly and the final HiRise scaffolds**. Each curve shows the fraction of the total length of the assembly in scaffolds of a given length or smaller. The fraction of the assembly is indicated on the Y-axis and the scaffold length in basepairs is given on the X-axis. The two dashed lines mark the N50 and N90 lengths of each assembly. This plot excludes scaffolds less that 1 kb.

**Table S1. Details of the draft assembly input and orientation into the final HiRise assembly**

**Fig S2. HiRise assembly improves the scaffold N50: 539 draft assembly scaffolds are reduced to 33**. Each bar represents a HiRise scaffold. Each color within the bar represents a draft assembly scaffold. Note: colors are used to show placement of input scaffolds but are not specific to any one scaffold. Number of draft input scaffolds are listed on right.

**Table S2. Summary mapping results for each ddRADseq library**

**Fig S3. Variant calling pipeline and results**. (A) The Venn diagram (top) shows the number of unique and shared variants detected by Platypus, Sentieon and SamTools variant callers. The flow chart beneath outlines the filtering strategy and the number of retained variants after each filtering step. (B) For the 884,092 variants detected by and passing filter flags in each variant caller, plot shows the mean proportion of variant calls that were concordant (Concord), discordant (Discord) or Missing between Sentieon and Platypus (Plat) and Sentieon and SamTools (SamT) among all samples as a function of coverage. Blue box highlights criterion range of coverage. Blue box highlights criterion range of. ≤65x coverage. (C) The observed versus expected heterozygosity ratio of each variant, represented as an open circle and plotted by minor allele frequency (MAF). Those with a ratio >1.2, above the black horizontal line, were filtered from the dataset. 902 Each variant is colored according to its Hardy-Weinberg equilibrium test statistic p-value bin listed in the legend (top left of plot). (D) For the 575,178 variants passing all filtering steps, the plot shows the mean proportion of variant calls that were concordant (Concord), discordant (Discord) or Missing between duplicate libraries of 4 samples (two samples from UW OshKosh, WI, “OK.TR1” and “OK.TR2”; and two samples from IL, “IL.TR1” and “IL.TR2”) as a function of coverage.

**Fig S4. ADMIXTURE 5-fold cross-validation (CV) error for each value of *K***. Shown are the CV error values for *K*=2 through *K*=8.

**Table S3. Pairwise FST estimates for the *K*=6 ADMIXTURE populations**
See text for labeling.

**Fig S5. Principal components PC17 and PC19 of all 153 genotyped squirrels reveal population structure within the OshKosh subset of squirrels**. Coloring is the same as in Fig 2A and 2C.

**Table S4: Biological and environmental data for each Oshkosh squirrel in which torpor onset was recorded**

**Table S5. Heart *trans*-eQTL results**

Table lists results for all *trans*-eqtl associations with p≤1×10^−5^

See Table 3 for SNP #

Tag ID is from the original Edge-tag datasets in [48,49]

β is effect size estimated from quantile-normalized values

**Table S6. SkM *trans*-eQTL results**

Labeling is the same as in Table S5

**Table S7. Liver *trans*-eQTL results**

Labeling is the same as in Table S5

**Table S8. BAT *trans*-eQTL results**

Labeling is the same as in Table S5

